# *Tudor-SN* promotes early replication of dengue virus in the *Aedes aegypti* midgut

**DOI:** 10.1101/751362

**Authors:** Sarah Hélène Merkling, Vincent Raquin, Stéphanie Dabo, Isabelle Moltini-Conclois, Lionel Frangeul, Hugo Varet, Maria-Carla Saleh, Louis Lambrechts

## Abstract

Diseases caused by mosquito-borne viruses have been on the rise for the last decades, despite the implementation of vector control methods primarily based on insecticides. An alternative control method currently in development is the use of lab-engineered mosquitoes that are incapable to carry viruses. This has stimulated efforts to identify optimal target genes that are naturally involved in mosquito antiviral defenses or required for viral replication. Although several antiviral immune pathways such as RNA interference (RNAi) have been previously characterized in mosquitoes, the genes that prevent or promote early viral replication in the midgut remain elusive. Here, we investigated the role of a member of the Tudor protein family, Tudor-SN, upon dengue virus infection in the mosquito *Aedes aegypti*. *Tudor-SN* expression was upregulated early after dengue virus infection and was subsequently positively correlated with viral loads in the midgut. Using RNAi-mediated knockdown, we showed that the loss of Tudor-SN reduced dengue virus replication in the *Ae. aegypti* derived cell line Aag2 and in the midgut of *Ae. aegypti* females *in vivo*. Using immunofluorescence assays, we found that *Tudor-SN* localizes to the nucleolus in both *Ae. aegypti* and *Aedes albopictus* cells. Finally, we used a reporter assay to demonstrate that Tudor-SN was not required for RNAi function *in vivo*. Collectively, these results define a novel proviral role for Tudor-SN upon early dengue virus infection of the *Ae. aegypti* midgut.

## Introduction

The mosquito *Aedes aegypti* transmits a wide range of pathogens to humans, many with severe consequences on public health, including dengue, Zika, and chikungunya viruses (1). For instance, dengue virus (DENV) infects 390 million people annually (2) and 50% of the world’s population is at risk for infection (3). DENV belongs to the *Flaviviridae* family and has a positive-sense, single-stranded RNA genome. DENV exists as four genetic types (DENV-1, -2, -3 and 4) that are phylogenetically related and loosely antigenically distinct (4). In the wild, mosquitoes acquire DENV by feeding on a viremic host. After the infectious blood meal, DENV infection is first established in the mosquito midgut before spreading systematically and reaching the salivary glands, where the virus engages in further replication (5) before being transmitted to the next host via the saliva released during the bite (6, 7).

The primary prevention strategy against arboviral diseases relies on the control of vector populations. Current vector control methods are mainly based on insecticides. Despite having been applied for decades, the burden of arboviral diseases keeps increasing (8). Human travel, urbanization, climate change, and geographic expansion of mosquito vectors increase pathogen transmission and spread (9). Over the last two decades, research efforts have led to the production of lab-engineered mosquitoes that either suppress wild vector populations or render them incapable of transmitting pathogens (10, 11). As the methods for genetic modification of mosquitoes develop, the need to identify optimal target genes that are naturally involved in mosquito antiviral defenses or required for viral replication also increases. Preferably, such pro- or antiviral target genes would act early during the course of an infection, and, when engineered, would permit early blocking of virus replication, at the level of the midgut cells. This would hinder viral dissemination and make further transmission of the virus impossible.

The majority of our knowledge about insect antiviral immunity originates from investigations in the model organism *Drosophila melanogaster* (12, 13), while studies in mosquito vectors remain more limited (14–16). The Toll, IMD, and Jak-Stat pathways have been implicated in insect innate immune responses to bacteria, fungi, viruses, and parasites. Their activation triggers translocations of NK-ΚB-like or Stat transcription factors to the nucleus, inducing the expression of an array of immune genes encoding antimicrobial peptides and virus restriction factors, amongst others (12–16). Another major branch of insect innate immunity is RNA interference (RNAi) which encompasses several pathways leading to the production of small RNA molecules of different characteristics, such as small interfering RNAs (siRNAs), microRNAs (miRNAs), and P element-induced wimpy testis (PIWI)-interacting RNAs (piRNAs) (17). The siRNA pathway is hitherto considered as the cornerstone of antiviral immunity in insects. It is initiated with the sensing and cleavage of viral double-stranded RNA (dsRNA) into 21-nucleotide-long siRNAs by the endonuclease Dicer-2. These siRNAs are loaded in the RNA-induced silencing complex (RISC) that guides Ago2-mediated cleavage of viral target sequences (17). Numerous studies reported that depletion of siRNA pathway components in mosquitoes resulted in increased arbovirus replication (18–22).

Although several pathways involved in antiviral immunity have been characterized in mosquitoes, several aspects of anti-DENV defense remain elusive. For example, the siRNA pathway was shown to inefficiently restrict DENV replication in the *Ae. aegypti* midgut (23). Besides, most of previous studies have focused on mosquito antiviral or restriction factors that antagonize DENV, but little is known about mosquito host factors with a proviral function, that is factors enhancing DENV propagation. Several human factors required for DENV infectivity were recently discovered through genome-wide CRISPR screens (24–26), whereas only a handful of DENV host factors have been identified in mosquitoes to date (27–30). Although CRISPR screens cannot be readily carried out in live mosquitoes, transcriptome analysis by high-throughput RNA sequencing is a powerful method to identify DENV host and restriction factors *in vivo* (31). For example, novel DENV restrictions factors (DVRF-1 and -2) that depend on the Jak-Stat pathway activation have been uncovered by overlapping transcriptional profiles of DENV-infected mosquitoes and mosquitoes with a hyperactive Jak-Stat pathway (32).

In this study, we exploited a unique transcriptomic dataset that we previously generated by performing RNA sequencing on individual midguts in a field-derived *Ae. aegypti* population during early DENV-1 infection (30). In addition to a conventional pairwise comparison of gene expression between DENV-infected and uninfected controls, we also used an approach to detect correlations between viral RNA load and gene expression. Out of 269 candidate genes identified by either method, only four were differentially expressed upon DENV infection and had expression levels that correlated with viral RNA load in infected mosquitoes (30). Amongst the four candidate genes identified by both methods was a gene encoding a member of the Tudor protein family, Tudor Staphylococcal Nuclease (abbreviated Tudor-SN or TSN), which we selected for further investigation in the present study. Using RNAi-mediated gene knockdown *in vivo*, we found that reduced *TSN* expression resulted in lower viral loads *in vitro* and *in vivo*. Immunofluorescence assays revealed that TSN localized to the nucleolus, and did not colocalize to DENV replication sites in DENV infected cells. Finally, we demonstrated that TSN was not involved in RNAi function in the midgut of adult mosquitoes. Altogether, our results demonstrate that TSN has an early proviral effect on DENV replication in the midgut and could be considered as a target to develop genetically-modified mosquitoes that are refractory to DENV infection.

## Results

### TSN *expression is upregulated upon DENV infection and positively correlates with viral loads*

Our previous transcriptomic analysis revealed that *TSN* (*AAEL000293)* expression was significantly upregulated upon DENV infection relative to mock controls one day after exposure to the infectious blood meal (Figure 1A), but not four days post blood meal (Figure 1B). Inversely, we found that *TSN* expression was not significantly correlated with DENV viral loads one day post blood meal (Figure 1C), but was positively correlated with DENV RNA loads four days post blood meal (Figure 1D). Thus, we concluded that *TSN* expression was induced by DENV infection within 24 hours after the infectious blood meal, and that subsequently, its expression was positively correlated with DENV replication. The positive correlation was suggestive of a proviral role for *TSN* upon DENV infection.

**Figure 1.**
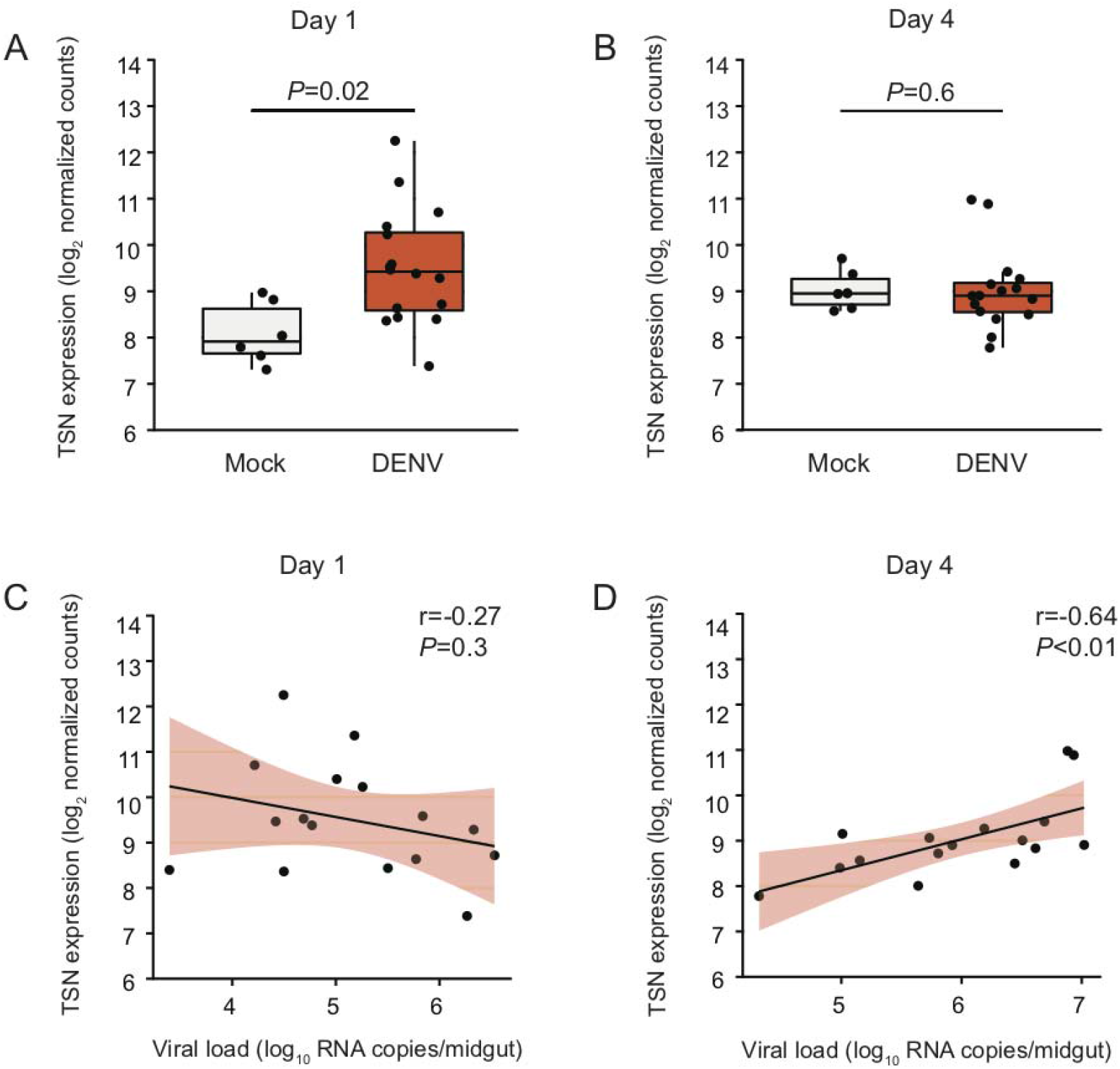
*TSN* is upregulated upon DENV infection and correlates positively with midgut viral loads. (**A**) *TSN* midgut expression levels on day 1 and day 4 post DENV exposure. Log_2_-transformed *TSN* normalized RNA-Seq counts are shown in mock-infected (n = 6) and DENV-infected (n = 16) midguts. *P* values of the pairwise t-tests are indicated. (**B**) Correlation of *TSN* expression level and viral load in DENV-infected midguts on day 1 and day 4 post virus exposure. Log_2_-transformed *TSN* normalized RNA-Seq counts are shown as a function of the log_10_-transformed midgut viral load. Black lines represent the linear regression and light purple shaded areas represent the 95% confidence intervals of the regression. Pearson’s coefficients of determination (r) and *P* values of the linear regression coefficient are indicated.

### *TSN is a DENV proviral factor* in vitro

First, we sought to test the proviral role of TSN *in vitro* by using *Ae. aegypti* Aag2 cells in culture. We transfected Aag2 cells with dsRNA to trigger RNAi-mediated knockdown of *TSN* and *GFP* and subsequently inoculated them with DENV at a multiplicity of infection of 1. We measured *TSN* expression levels by reverse transcription quantitative PCR (RT-qPCR) at 0, 12, 24, 36, 48, 72 and 96 hours post infection, and found that *TSN* knockdown efficiency ranged from ≈50% to 80% and was statistically significant at most of the time points (Figure 2A). We visualized TSN protein levels by Western blotting using an antibody directed against the human ortholog of TSN named SND1, which also reacted with the *Ae. aegypti* TSN. We confirmed that *TSN* knockdown reduced TSN protein levels by 70 to 80% in Aag2 cells at 24 and 48 hours post DENV infection, compared to the GFP control (Figure 2B). The knockdown effect was strongest at 24 hours post infection, most likely because the transfection has a transient effect on *TSN* expression that diminishes over time. To determine whether TSN also augmented viral infection *in vitro*, were measured both DENV RNA levels (Figure 2C) and DENV infectious titers by focus-forming assay (Figure 2D) over the course of infection. We found that DENV RNA levels were significantly reduced upon *TSN* knockdown relative to control levels at 24 hours post infection (Figure 2C, *P*<0.01). Moreover, DENV RNA levels were consistently lower in *TSN*-depleted cells from 24 to 96 hours post infection. DENV infectious titers were also significantly reduced upon *TSN* knockdown at 24 and 48 hours post infection (Figure 2D, *P*<0.05 and *P*<0.001, respectively). Overall, these data demonstrated a proviral role of TSN *in vitro*.

**Figure 2.**
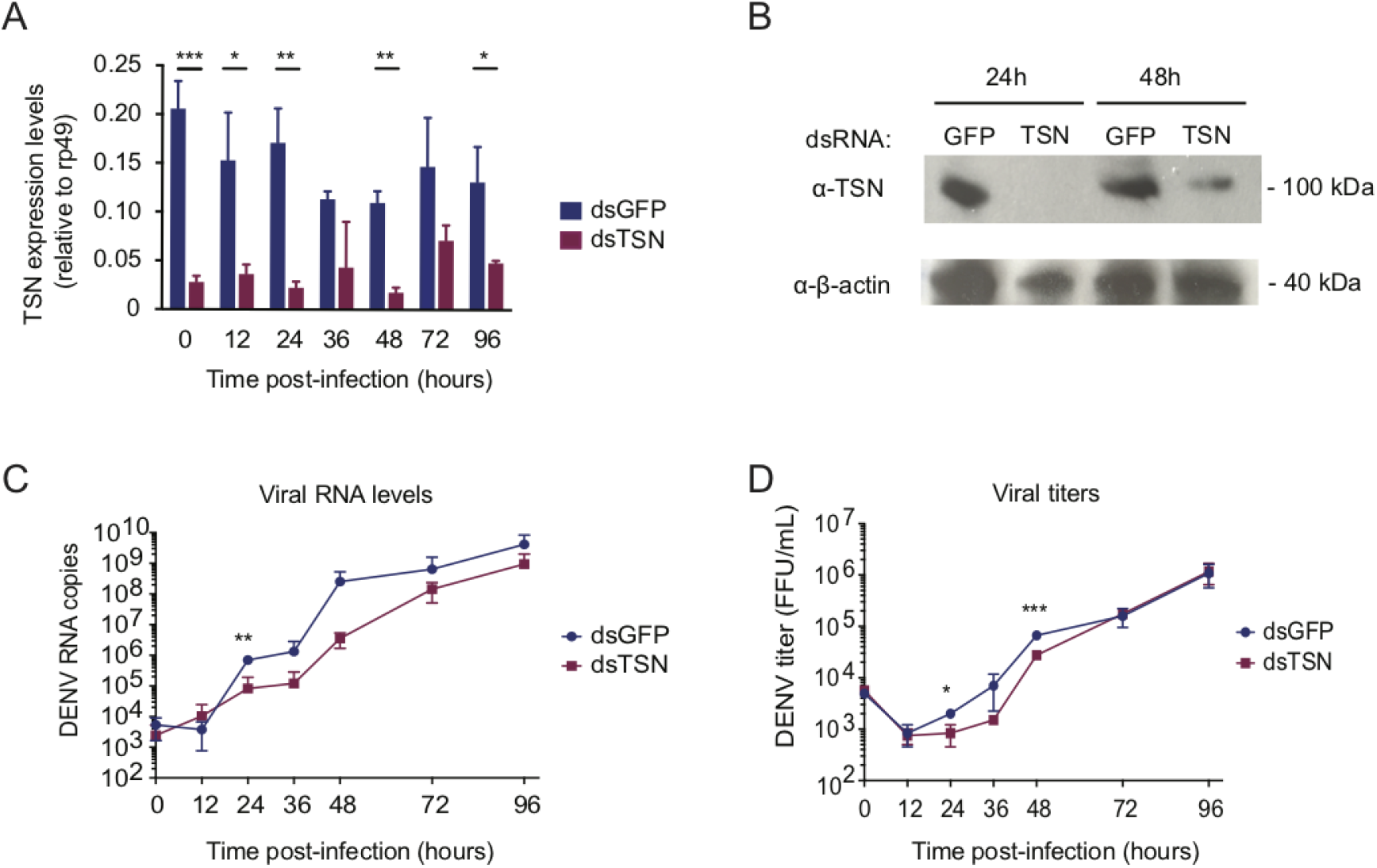
*TSN* promotes DENV infection in *Ae. aegypti* Aag2 cells. (**A**) *TSN* expression levels in Aag2 cells transfected with dsRNA targeting *TSN* (dsTSN) or *GFP* (dsGFP), over the course of DENV infection at a multiplicity of infection of 0.1. Aag2 cells were transfected 3 and 1 day prior to virus infection. *TSN* expression was measured by RT-qPCR and normalized to *rp49*. Data represent mean and standard deviation of three biological replicates. \**P* < 0.05; \*\**P* < 0.01; \*\*\**P* < 0.001 (Student’s t-test). Data is from the same experiment shown in panels C and D. (**B**) Western blot analysis of *TSN* expression in Aag2 cells 24 and 48 hours after viral infection. An anti-β-actin monoclonal antibody was used for loading control. Molecular mass in expressed in kilo Daltons (kDa). (**C**, **D**) Analysis of DENV RNA levels (**C**) or DENV infectious titers (**D**) over the course of DENV infection following *TSN* or *GFP* knockdown. RNA levels were measured by RT-qPCR on RNA from cell extracts, and infectious titers were determined by focus-forming assay on cell culture supernatants. Data represent mean and standard deviation of three biological replicates. \**P* < 0.05; \*\**P* < 0.01; \*\*\**P* < 0.001 (Student’s t-test).

### TSN *is a DENV proviral factor* in vivo

To confirm the proviral role of *TSN in vivo*, we experimentally reduced *TSN* expression in adult female mosquitoes by intrathoracic injection of dsRNA and subsequently exposed them to an infectious blood meal containing 10^7^ focus-forming units (FFU)/mL of DENV (Figure 3A). First, we monitored *TSN* expression levels in individual mosquitoes by RT-qPCR on days 0, 1 and 4 after the blood meal. On day 0, which corresponds to three days after injection of the dsTSN, *TSN* expression was significantly knocked down relative to mosquitoes injected with a control dsRNA targeting an exogenous Green Fluorescent Protein (GFP) sequence (Figure 3B, *P*<0.001). Reduced *TSN* expression persisted over time through day 1 (Figure 3C, *P*<0.0001) and day 4 (Figure 3D, *P*=0.01) after exposure to the infectious blood meal. Importantly, reduced *TSN* expression did not significantly impact the survival of mosquitoes during the seven days following injection compared to the GFP control (Figure 3E, *P*=0.54). Next, we measured DENV RNA loads by RT-qPCR and found a ≈50% reduction of viral loads in mosquito midguts depleted for *TSN*, compared to the GFP control, four days after the infectious blood meal (Figure 3F, *P*<0.0001). We also measured DENV RNA loads one day after the infectious blood meal and did not observe a decrease of viral loads. However, this is most likely due to the presence of viral RNA in the undigested blood still present in the midgut at this time point, as shown in a previous report (30). These results confirmed the proviral role of *TSN* during DENV infection in the mosquito midgut four day after the infectious blood meal. Finally, we asked whether *TSN* knockdown had an impact on infection prevalence after DENV exposure. Amongst the mosquito midguts analyzed by RT-PCR on day 4, we found that 88% and 90% were positive for DENV RNA in the *TSN* knockdown and the dsGFP control groups, respectively (*P*=0.59). Overall, our data demonstrated that although *TSN* does not influence the probability of DENV infection, it promotes early DENV replication in the mosquito midgut.

**Figure 3.**
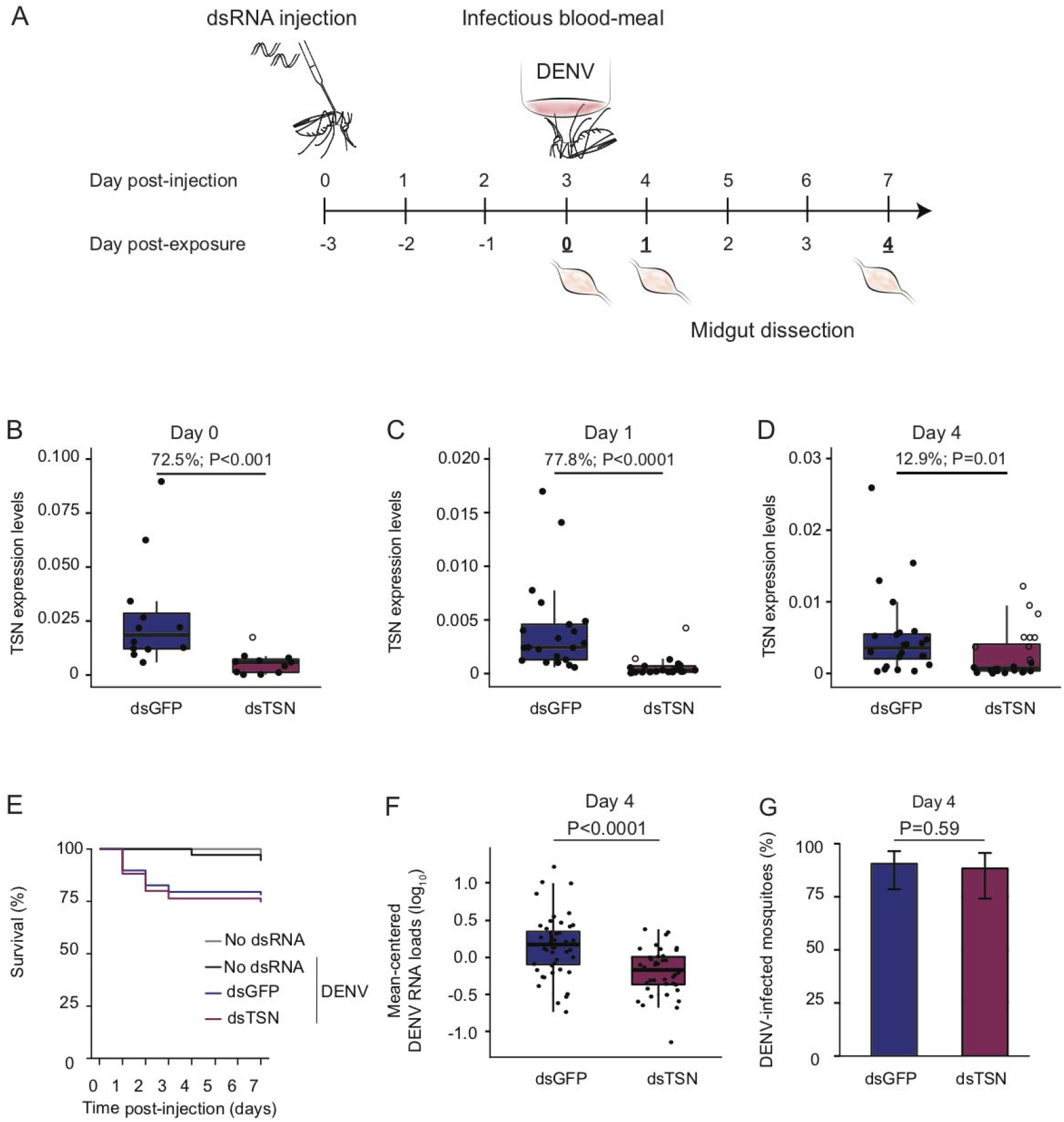
*TSN* promotes DENV infection the mosquito midgut. (**A**) Experimental scheme of the gene-silencing assays *in vivo*. (**B**, **C**, **D**) *TSN* expression levels following gene knockdown on day 0 (**B**), day 1 (**C**) and day 4 (**D**) after exposure to DENV infectious blood meal. Mean percentage of gene expression knockdown at day 0 (**B**), day 1 (**C**) and day 4 (**D**) after DENV exposure are indicated. Boxplots show *TSN* expression normalized by *rp49* and expressed as 2^-dCt^ values in n = 12-24 individual mosquito midguts per group. Individuals with less than 50% gene expression knockdown are shown as empty dots. Data are representative of three separate experiments. *P* values above the graph indicate statistical significance assessed with a Wilcoxon test. (**E**) Percentage of survival following dsRNA injection and/or DENV exposure. Mosquitoes were injected with dsRNA targeting *TSN* (n = 127) or targeting *GFP* (dsGFP, n = 110) 3 days prior to DENV exposure. Mosquitoes non injected (non inj., n = 71) or fed on a non-infected blood meal (mock, n = 62) were used as controls. No significant difference in mortality was detected between dsGFP and dsTSN mosquitoes according to a Cox model (*P* = 0.54). (**F**) DENV RNA levels in mosquito midguts dissected from mosquitoes previously injected with dsGFP (n = 53) or dsTSN (n = 55). Boxplots represent the viral load measured by RT-qPCR on day 4 post exposure. Data represent three separate experiments combined. Viral loads are adjusted for differences between experiments and expressed in mean-centered DENV RNA loads. The *P* value above the graph indicates statistical significance of the treatment effect assessed with an analysis of variance accounting for the experiment effect. (**G**) DENV infection prevalence in mosquito midguts measured by RT-qPCR on day 4 post virus exposure following injection with dsGFP (n = 53) and dsTSN (n = 43). Data from three separate experiments were combined after verifying the lack of a detectable experiment effect. Error bars represent 95% confidence intervals of the percentages. The *P* value above the graph indicates statistical significance of the treatment effect assessed with a logistic regression.

### TSN localizes to the nucleolus in mosquito cells

It was previously shown that TSN could interact with DENV RNA in mammalian cells (33), which lead us to ask whether TSN co-localized with DENV-derived RNA in mosquito cells. Double-stranded RNA is produced during the replication of single-stranded RNA viruses like DENV and is a hallmark of virus infection (34). Previous reports demonstrated that antibodies directed against dsRNA did not cross-react with cellular rRNA or tRNA and could be used to identify flavivirus replication complexes in infected cells (35). To determine the subcellular localization of TSN, we performed immunofluorescence assays in mosquito cells derived from *Ae. albopictus* (C6/36, Figure 4A) or *Ae. aegypti* (Aag2, Figure 4B) using the anti-SDN1 antibody previously validated by Western blotting (Figure 3B) and a monoclonal antibody targeting dsRNA (called αK1). In both cell types, we found that TSN was expressed and localized to the nucleolus. Indeed, it localized to the nucleus region but did not overlap with DAPI staining, which is reported to exclude the nucleolus (36). Moreover, the staining was more intense at the nucleus-nucleolus interface where it formed a “ring”. TSN localization did not change upon DENV infection, nor did its expression level. Six days after DENV infection of C6/36 and Aag2 cells, dsRNA staining was readily detectable and mainly localized to cytoplasmic regions of infected cells, likely corresponding to viral replication sites. Since TSN localized to the nucleolus, and the dsRNA to the cytoplasm, we did not observe overlapping signals between both stainings. Thus, we conclude that TSN does not interact with DENV RNA at its replication site. However, it remains possible that interactions occur with other forms of DENV RNA (positive or negative single-stranded RNA), or viral proteins.

**Figure 4.**
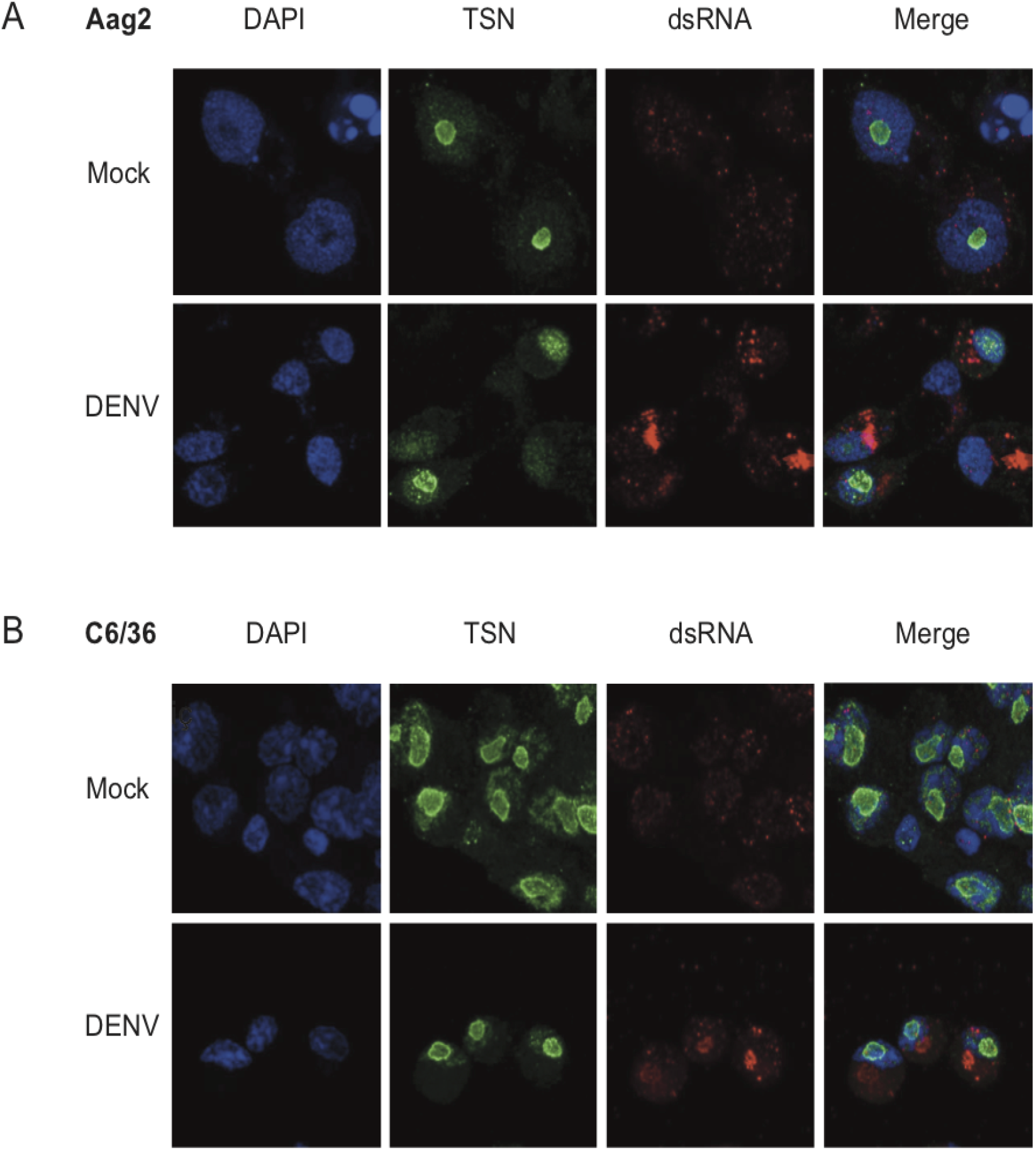
*TSN* localizes to the nucleolus in *Ae. aegypti* and *Ae. albopictus* cells. (**A**, **B**) Immunofluorescence assays on *Ae. aegypti* Aag2 (**A**) or *Ae. albopictus* C6/36 (**B**) cells 6 days after infection with DENV. Cells were stained with the nuclear stain DAPI, an antibody against TSN coupled to an Alexa-488 secondary fluorescent antibody, and an antibody against dsRNA (αK1) coupled to an Alexa-689 secondary fluorescent antibody. Cells were imaged on a confocal microscope at 63X magnification.

### *RNAi is functional in* TSN-*depleted mosquitoes*

Proteins containing Tudor motifs have been implicated in multiple aspects of RNA metabolism such as RNA splicing or small RNA pathways (37, 38). TSN was shown to be a component of the RISC in *Caenorhabditis elegans*, *Drosophila* and mammals (39) and was suggested to participate in RNAi function in the tick *Ixodes scapularis* (40). Therefore, we asked whether TSN was involved in RNAi function in *Ae. aegypti*. We adapted a luciferase-based RNAi sensor assay developed in *Drosophila* to mosquitoes (41–43). Adult females were intrathoracically injected with a mix of lipofectant along with Firefly *luciferase* reporter plasmid with Firefly *luciferase*-specific dsRNA and dsRNA targeting *GFP* (as a control), *TSN* or *Ago2* (Figure 5A). A reporter plasmid encoding a Renilla *luciferase* was used as an *in vivo* transfection control. Three days after injection, the efficiency of Firefly *luciferase* silencing was measured in whole mosquito homogenates (Figure 5B). When reporter plasmids were injected together with control dsGFP, we observed a wide range of luminescence counts (likely due to variable *in vivo* transfection efficiency), but the average luciferase activity was about 100-fold higher than when dsRNA targeting Firefly *luciferase* (dsLuc) was co-transfected with the reporter plasmids and control dsGFP. The silencing of Firefly *luciferase* was restored upon knockdown of *Ago2*, a key component of the RNAi pathway, demonstrating the validity of the reporter assay. Finally, we observed that the silencing of Firefly *luciferase* was maintained upon co-transfection with the dsRNA targeting *TSN*, suggesting that *TSN* is not involved in RNAi function in *Ae. aegypti* (Figure 5B). We measured expression levels of *TSN* and *Ago2* upon co-transfection with reporter plasmids and dsRNA, and verified that *TSN* and *Ago2* expression levels were significantly reduced upon knockdown with their specific dsRNA (Figure 5C and 5D). Although we cannot exclude that residual *TSN* expression could suffice to maintain its activity, the knockdown efficiency was similar to that of *Ago2*. Overall, these results support the conclusion that *TSN* is not essential for RNAi function in *Ae. aegypti*.

**Figure 5.**
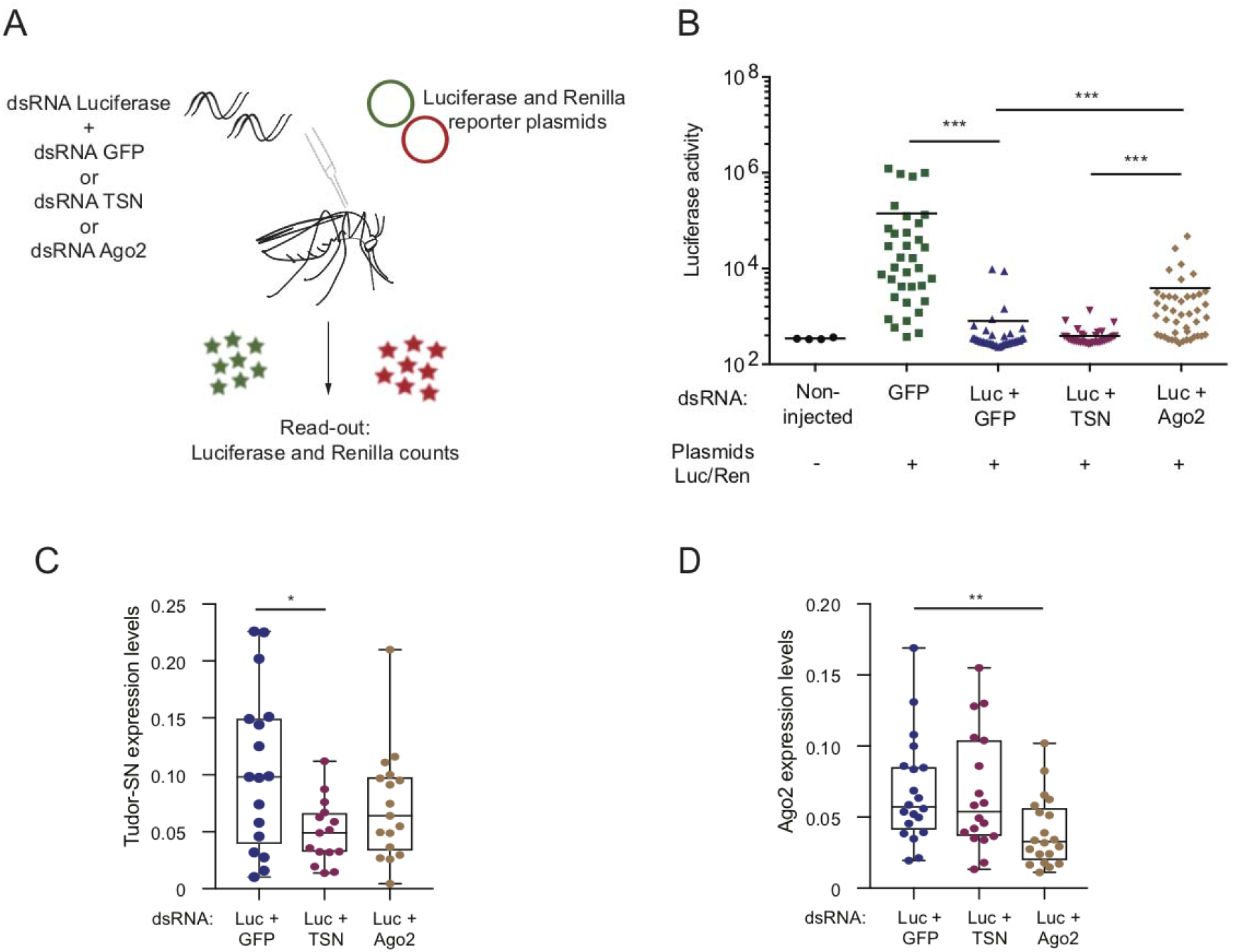
*TSN* is not required for RNAi function. (**A**) Experimental scheme. Adult mosquitoes were co-injected with Firefly *luciferase* (Fluc) and Renilla *luciferase* (Rluc) reporter plasmids, and dsRNA targeting Fluc in combination with dsRNA against GFP, TSN or Ago2. Luminescence was measured 3 days post injection. (**B**) *In vivo* RNAi reporter assay. Reporter gene activity was measured in individual mosquitoes. Luminescence counts of Firefly luciferase are shown as absolute values. Mean and standard deviation are shown. \**P* < 0.05; \*\**P* < 0.01; \*\*\**P* < 0.001 (Mann-Whitney t-test). (**C**, **D**) Expression levels of *TSN* (**C**) and *Ago2* (**D**) measured by RT-qPCR on mosquitoes harvested 3 days after injection with plasmids and dsRNA. Gene expression was normalized by *rp49* and expressed as 2^-dCt^ values. \**P* < 0.05; \*\**P* < 0.01; \*\*\**P* < 0.001 (Student’s t-test). Data is from the same experiment shown in panel B.

## Discussion

Antiviral immunity in *Ae. aegypti* mosquitoes remain poorly understood. Using a novel approach of transcriptomic analysis, we previously uncovered four genes that not only responded to DENV infection in the mosquito midgut but also had expression levels that correlated with viral loads in infected mosquitoes (30). Here, we focused on one of these four genes, *Tudor-SN*, encoding a member of the Tudor protein family. *Tudor-SN* was induced upon DENV infection, and its expression correlated positively with viral RNA load (30). Using RNAi-mediated knockdown assays *in vivo* and *in vitro*, we demonstrated that *TSN* promotes DENV replication. We performed localization studies and discovered that TSN localizes to the nucleolus of *Ae. aegypti* and *Ae. albopictus* cells, and does not colocalize with DENV replication sites. Moreover, we found that despite belonging to the Tudor family, TSN was not essential for RNAi function in adult mosquitoes.

TSN is a known component of the RISC, the RNAi protein complex that carries siRNAs and directs cleavage of complementary viral sequences in *Caenorhabditis elegans*, *Drosophila* and mammals (39). Additionally, previous work in *Drosophila* and other model organisms found essential functions for Tudor domain-containing proteins in piRNA biogenesis. A recent study describing a functional knockdown screen of all predicted *Ae. aegypti* Tudor proteins did not reveal a role for Tudor-SN in piRNA biogenesis (44). This finding is consistent with our observation that Tudor-SN is not necessary for RNAi function *in vivo* in *Ae. aegypti*. Finally, Tudor-SN was also shown to be a conserved component of the basic RNAi machinery in *Ixodes* ticks (40). This study reported an effect of Tudor-SN on dsRNA-mediated gene silencing, which possibly involves the siRNA pathway. However, no evidence was obtained for a role of Tudor-SN in the response to microbial infection (40). Taken together, the data described in this study and discussed above did not find strong links between RNAi function and Tudor-SN in arthropods. Further studies using knockout mutants might be necessary to confirm these findings.

The mammalian ortholog of Tudor-SN is generally referred to as p100 and was identified as a host factor interacting with the 3’ untranslated region of the DENV genome (45). Moreover, *p100* knockdown led to reduced levels of viral RNA and protein in mammalian cells, providing evidence that p100 was required for efficient DENV replication. Although these results are consistent with those reported here, in mammalian cells p100 was shown to interact with DENV genomic RNA and dsRNA replication intermediates, which we did not observe. Importantly, the subcellular localization of p100 in mammalian cells was perinuclear, whereas it was nucleolar in mosquito cells. This discrepancy in localization hints to a divergence in function between mammals, in which p100 interacts directly with viral RNA, and insects, for which evidence is lacking. Microscopy-based localization studies being limited in sensitivity and resolution, elucidating Tudor-SN function in mosquitoes will require further experiments to more definitely exclude interactions between viral RNA and Tudor-SN, such as protein immunoprecipitation and sequencing of associated RNA (RIP-seq).

The predicted structure of *Ae. aegypti* Tudor-SN includes four Staphylococcal Nuclease (SN)-like domains and a Tudor domain embedded in a fifth SN domain. Tudor-SN homologs are found in diverse eukaryotic species such as plants, humans (Staphylococcal Nuclease and Tudor domain containing 1) and insects (*Drosophila* Tudor-SN). The very similar structure of eukaryotic Tudor-SN homologs is consistent with potentially conserved functions (Figure 6). However, Tudor-SN subcellular localization is variable in other species, which also hints towards species-specific function(s) of Tudor-SN proteins.

**Figure 6.**
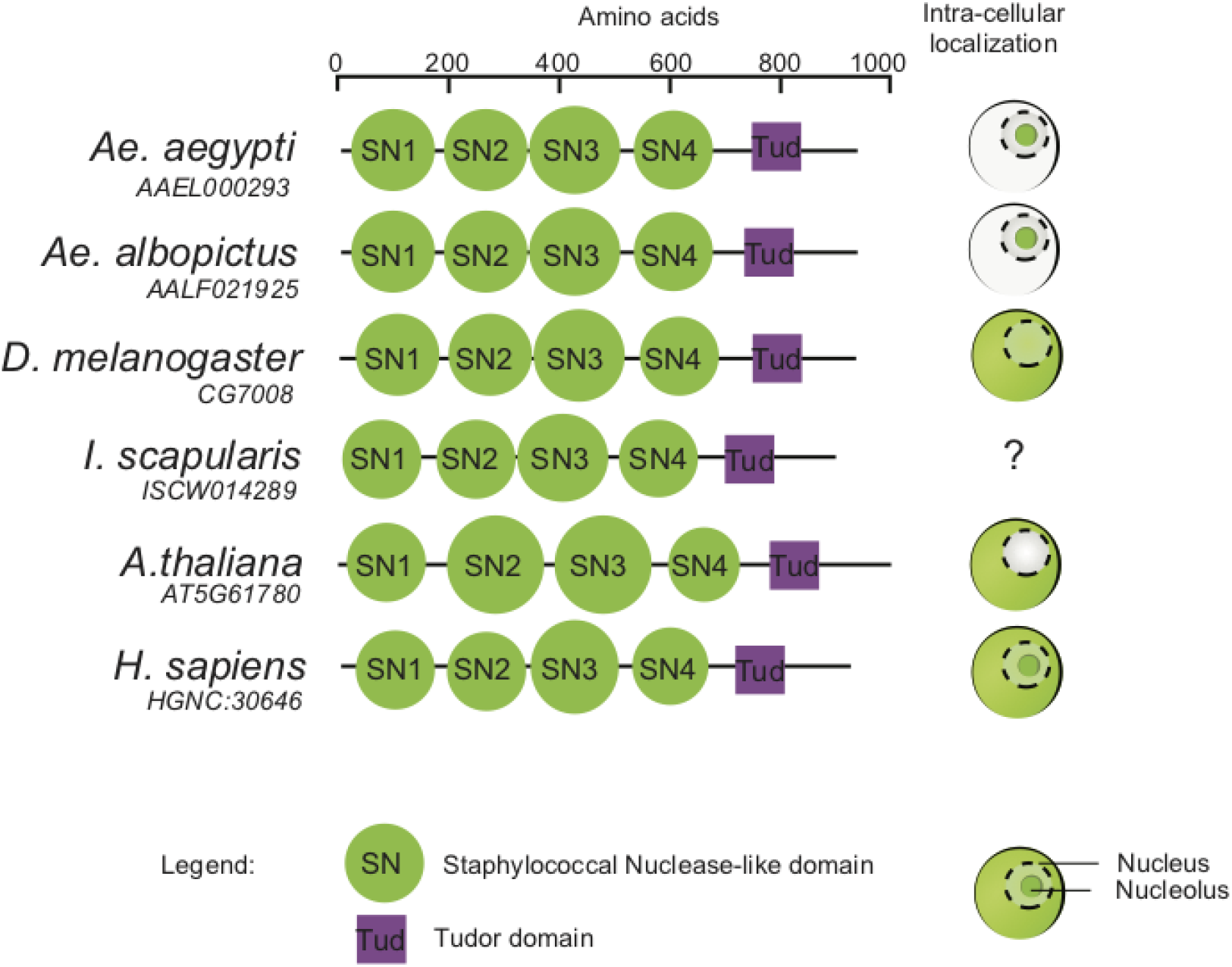
Conserved structure of TSN homologs among diverse eukaryotic species. Structural domains provided in the UniProt database are shown for homologs of *Ae. aegypti* TSN (Q17PM3) in *Drosophila melanogaster* (Q9W0S7), *Ixodes scapularis* (B7QIP4), *Arabidopsis thaliana* (F4K6N0) and *Homo sapiens* (Q7KZF4). Diagrams on the right side indicate TSN subcellular localization (in red) in the cytoplasm, nucleus and/or nucleolus of each species based on the literature (39, 62–65).

Our observation that *Ae. aegypti* Tudor-SN localizes primarily to the nucleolus of mosquito cells makes it unlikely that its proviral effect on DENV relies on a direct action on viral genome stability or replication. The nucleolus is a multifunctional nuclear domain involved in ribosome biogenesis and several other cellular functions, such as cell cycle regulation, telomere metabolism or DNA damage sensing and repair (46). Various nucleolar alterations during viral infection have been documented (47). Interestingly, DENV non-structural protein 5 (NS5), which encodes the virus RNA-dependent RNA polymerase, was recently shown to localize to the nucleolus of infected mammalian cells (48), where it interferes with precursor messenger RNAs (pre-mRNA) splicing to limit host antiviral response (49). Human Tudor-SN was implicated in spliceosome assembly and therefore, may influence splicing of pre-mRNAs and/or interact with DENV NS5 to facilitate viral RNA accumulation (50). More generally, Tudor-SN could promote viral replication through regulation of gene expression. For example, the Jak-Stat pathway protects *Ae. aegypti* against DENV infection (32) and Tudor-SN was showed to bind Stat proteins to modulate host gene transcription (51). Also, Tudor-SN is a component of stress granules (52) and could interfere with their formation to facilitate DENV infection (53). Particularly interesting is the early proviral effect of *TSN* in the mosquito midgut, which might suggest a role for *TSN* in viral sensing, or early antiviral responses. For instance, *TSN* might sense the infection and, in the nucleus, alter the spliceosome to increase the availability of cellular resources that the virus requires to replicate. Its presence in the nucleolus and at the nucleus-nucleolus interface might enhance ribosome biogenesis and subsequently increase production of viral proteins.

Although several pathways involved in antiviral immunity have been characterized in mosquitoes, several aspects of anti-DENV defense remain to be elucidated, particularly during the early phase of infection. For example, the siRNA pathway was shown to inefficiently restrict DENV replication in the *Ae. aegypti* midgut (23). The present work adds to the small number of studies that identified DENV proviral factors in mosquitoes (27–30). Such host factors have been proposed as new targets for antiviral therapy in humans (24–26). Although the development of novel vector control methods has focused on viral restriction factors so far (54), targeting essential host factors could complement antiviral strategies in mosquitoes. We showed that *Tudor-SN* is such a host factor for DENV in the mosquito *Ae. aegypti*. Further studies will be necessary to elucidate the specific mechanisms underlying the role of Tudor-SN in DENV replication.

## Material and Methods

### Ethics

The Institut Pasteur animal facility has received accreditation from the French Ministry of Agriculture to perform experiments on live animals in compliance with the French and European regulations on care and protection of laboratory animals. This study was approved by the Institutional Animal Care and Use Committee at Institut Pasteur under protocol number 2015–0032.

### Cells and virus

C6/36 cells (derived from *Ae. albopictus*) and Aag2 cells (derived from *Ae. aegypti*) were cultured in Leibovitz’s L-15 medium (Life Technologies) supplemented with 10% fetal bovine serum (FBS, Life Technologies), 1% non-essential amino acids (Life Technologies) and 0.1% Penicillin-Streptomycin (Life Technologies) at 28°C. DENV type 1 (DENV-1) isolate KDH0030A was originally derived in 2010 from the serum of a dengue patient at the Kamphaeng Phet Provincial Hospital, Thailand (55). The full-length consensus genome sequence is available from GenBank under accession number HG316482. Virus stock was prepared in C6/36 cells and infectious titers measured in C6/36 cells using a standard focus-forming assay (FFA) as previously described (56).

### Mosquito rearing and experimental infections

All experiments were performed with adult *Ae. aegypti* mosquitoes derived from a field population originally sampled in 2013 in Thep Na Korn, Kamphaeng Phet Province, Thailand. Experiments took place within 16 generations of laboratory colonization. Mosquitoes were reared in standard insectary conditions, as previously reported (56). Experimental virus infections were performed in a level-3 containment facility, as previously described (56). Shortly, 5- to 7-day-old female mosquitoes were deprived of 10% sugar solution 24h before oral exposure to DENV. The infectious blood meal consisted of a 2:1 mix of washed rabbit erythrocytes and DENV-1 viral suspension (to reach an infectious titer of 10^7^ FFU/mL) supplemented with 10 mM ATP (Sigma). Mosquitoes were fed for 30 min through a pig-intestine membrane using an artificial feeder (Hemotek Ltd) set at 37°C. Fully engorged females were incubated at 28°C, 70% relative humidity and under a 12-hour light-dark cycle with permanent access to 10% sucrose till further use.

### RNA isolation from mosquito midguts

Midguts were dissected in 1x PBS, and immediately transferred to a tube containing 800 µL of Trizol (Life Technologies) and ~20 1-mm glass beads (BioSpec). Samples were homogenized for 30 sec at 6,000 rpm in a Precellys 24 grinder (Bertin Technologies). RNA was extracted as previously described (30), and stored at −80°C till further use.

### Reverse transcription and quantitative PCR

Viral RNA was reverse transcribed and quantified using a TaqMan based qPCR assay, using NS5-specific primers and 6-FAM/BHQ-1 double-labeled probe (sequences provided in Table 1). Reactions were performed with the Superscript III Platinum One-Step qRT-PCR kit (Life technologies) following the manufacturer’s instructions. *Tudor-SN* and *Ago2* expression levels were measured using a SybrGreen based qPCR assay, using gene-specific primers (sequences provided in Table 1). First, total RNA from individual midguts or adults was reverse transcribed into cDNA using MMLV reverse transcriptase (Invitrogen), according to manufacturer’s instructions. Quantitative PCR was performed on a LightCycler 96 real-time thermocycler (Roche) using SYBR Green MasterMix from Roche (Figure 1) or Promega (Figure and 4). qPCR efficiency and Ct values were unaffected by change of SYBR Green reagent. The qPCR programs was as follows: an initial denaturation step of 5 min at 95°C, followed by 40 cycles of 10 sec at 95°C, 20 sec at 60°C and 10 sec at 72°C. A melting curve was generated to confirm the absence of non-specific PCR amplicons using the following program: 5 sec at 95°C, 60 sec at 65°C and continuous fluorescence acquisition up to 97° C with a ramp rate 0.2°C/sec. Relative expression was calculated as 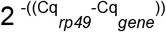, using the *Ae. aegypti* ribosomal protein-coding gene *rp49* (*AAEL003396*) for normalization.

**Table 1:**
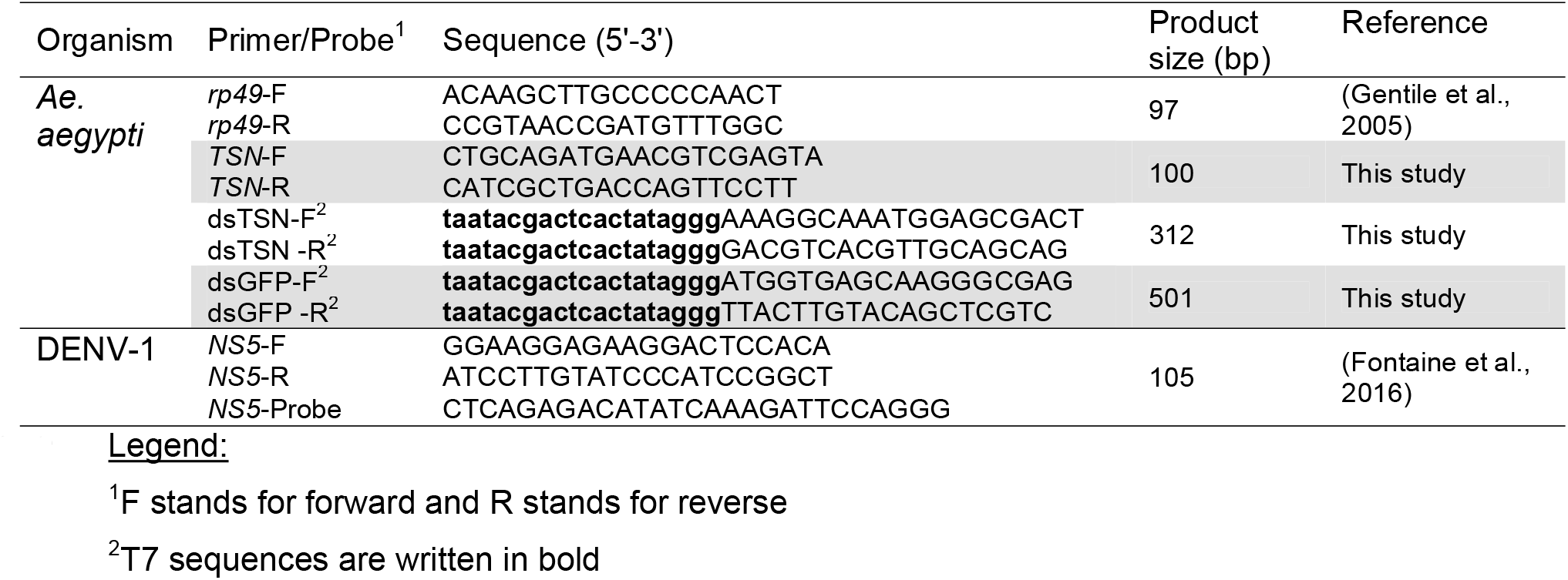
List of oligonucleotide primers and molecular probes used in this study

### Double-stranded RNA synthesis

Design and synthesis of dsRNA used in knockdown assays has been described previously (30). Briefly, dsRNA was synthesized from a GFP-containing plasmid or from a cDNA template produced by RT-PCR on RNA isolated from a pool of *Ae. aegypti* mosquitoes. T7 promotor sequences were incorporated by PCR to the amplicon that was used as a template for the synthesis using the MEGAscript RNAi kit (Life Technologies).

### Gene silencing assays *in vivo*

RNAi-mediated knockdown of target genes was performed as previously described (30). Briefly, 500 ng to 1 μg of dsRNA was injected intra-thoracically using Nanoject II or III (Drummond). Injection volume was 140 nL expect for the RNAi reporter assay described below. Control mosquitoes were injected with a dsRNA targeting Green Fluorescent Protein (GFP). After injection, mosquitoes were incubated for 3 days at 28°C before exposure to an infectious blood meal.

### RNAi reporter assay

RNAi competency of adult mosquitoes was assed using a reporter assay adapted from previously published methods in *Drosophila* (41–43). *In vivo* plasmid transfection method was optimized from protocols previously published for *Ae. aegypti* (57, 58). Five to seven-day-old adult mosquitoes were injected in the thorax using a Nanoject III (Drummond) with a suspension of ≈ 300 nL containing a 1:1 mixture of unsupplemented Leibovitz’s L-15 medium (Life technologies) and Cellfectin II (Thermo Fisher Scientific) complexed with 50 ng pUb-GL3 (encoding Firefly *luciferase*, FLuc), 50 ng pCMV-RLuc (encoding Renilla *luciferase*) described previously (59, 60), and 500 ng FLuc-specific and 1 μg *GFP*-, *TSN*- or *Ago2*-specific dsRNA. After incubation for 3 days at 28°C, mosquitoes were homogenized in passive lysis buffer (Promega) using the Precellys 24 grinder (Bertin Technologies) for 30 sec at 6,000 rpm. Samples were transferred to a 96-well plate and centrifugated for 5 min at 12000 x g. Fifty microliters of supernatant were transferred to a new plate, and 50 μL LARII reagent added for the first FLuc measurement. Next, 50 μL Stop&Glow reagent was added before the second measurement of RLuc, according to the Dual Luciferase assay reporter system (Promega). Counts of RLuc were used to control for transfection efficiency, and samples with less than 1,000 counts were discarded from the analysis. However, Renilla counts were not used for normalization due to great biological variation between samples. Data are presented as read counts for FLuc.

### Immunofluorescence assays

Mock- and DENV-infected Aag2 cells were fixed on coverslips using 4% paraformaldehyde (Sigma-Aldrich) for 30 min at room temperature (20-25°C). Following permeabilization with 1X PBS, 0.1% Triton-X100, cells were incubated with mouse anti-dsRNA αK1 (English & Scientific consulting) and rabbit anti-TSN antibodies diluted 1:500 in 1X PBS, 0.1% Triton-X100, 2% Normal Goat Serum for 1 hour at room temperature. Subsequently, cells were washed tree times with 1X PBS with 0.1% Triton X-100 and incubated with goat anti-mouse AlexaFluor 594 and goat anti-rabbit Alexa Fluor 488 diluted 1:1,000 in 1X PBS, 0.1% Triton-X100, 2% Normal Goat Serum (Life technologies), overnight at 4°C. After three washes in 1X PBS with 0.1% Triton X-100, cover slips were mounted on a glass slide in ~10 μL Prolong Gold anti-fade medium containing DAPI (Thermo Fisher) and imaged with a confocal microscope LSM 700 inverted (Zeiss) at 63X magnification.

### Gene silencing and DENV infection *in vitro*

Aag2 cells were transfected in 24-well plates with 500 ng of dsRNA using Lipofectamine LTX (Invitrogen) along with Plus reagent according to the manufacturer’s instructions. To increase knockdown efficiency, a second round of transfection with 500 ng of dsRNA was performed 48 hours after the initial transfection. Infection with DENV was performed 24 hours after the last transfection. Cells were incubated for 1 hour in L-15 infection medium containing 2% FBS and DENV at a multiplicity of infection of 1. After removal of the infectious inoculum, cells were refreshed with fully supplemented with L-15 medium and incubated at 28°C.

### Western blotting

Aag2 cells were harvested, washed once in PBS and resuspended in RIPA buffer (20 mM Hepes-KOH pH 7.5, 100 mM KCl, 5% glycerol, 0.05% NP40, with freshly added 0.1M DTT and complete, EDTA-free, protease inhibitors (Roche)). Cells were lysed in Laemelli buffer (Sigma-Aldrich) with 10% β-mercaptoethanol and incubated at 95°C for 5 min. Protein lysates were loaded on a 4–20% precast mini-protean polyacrylamide gel (Bio-Rad) then transferred to a nitrocellulose membrane. The blot was incubated for 1 hr at room temperature with rabbit anti-TSN antibody (Abcam 65078) diluted 1:200 in blocking buffer (5% skimmed milk powder, 0.1% Triton-X100 in 1X PBS). After 3 washes with 1X PBS, the blot was incubated with a secondary antibody, HRP-conjugated polyclonal goat anti-rabbit IgG (GE Healthcare) diluted 1:5,000 in 1X PBS, 0.1% Triton-X100 for 1 hour at room temperature. Next, the blot was incubated with a primary anti-β-actin murine antibody (Sigma-Aldrich, clone AC-74) at 1:6,000 dilution in blocking buffer, and an HRP-conjugated polyclonal goat anti-mouse IgG (GE Healthcare) diluted at 1:5,000 in 1X PBS, 0.1% Triton-X100 was used as a secondary antibody. Bound antibodies were revealed by chemiluminescence with the SuperSignal West Pico Chemiluminescent Substrate (Fisher Scientific). Quantification of band intensities was performed in ImageJ (61).

### Statistical analysis

Statistical analysis methods have been described previously (30). Additionally, statistical significance in figures 3 and 5 was tested using unpaired two-tailed Student’s t-test and Mann-Whitney test as implanted in GraphPad Prism version 7. *P* values below 0.05 were considered statistically significant.

### RNA-seq data availability

The RNA-Seq data were deposited to SRA under accession number PRJNA386455.

## Acknowledgements

We thank Lambrechts lab members and Marie-Agnès Dillies for helpful comments and discussions. We are grateful to Catherine Lallemand for assistance with mosquito rearing. We thank Alongkot Ponlawat for the initial field sampling of mosquitoes in Thailand. We thank Alain Kohl for sharing the reporter plasmids and Emilie Pondeville for her insights on the reporter assay protocol.

## Authors contributions

S.H.M., V.R., S.D., M.-C.S. and L.L. designed the experiments; S.H.M., V.R., I.M.-C. and S.D. performed the experiments; S.H.M., V.R., H.V., L.F. and L.L. analyzed the data; S.H.M., V.R., M.-C.S. and L.L. wrote the paper.

